# Drab and distant birds are studied less than their fancy-feathered friends

**DOI:** 10.1101/2023.10.26.560707

**Authors:** Silas E. Fischer, Joshua G. Otten, Andrea M. Lindsay, Donald B. Miles, Henry M. Streby

## Abstract

Human decisions are influenced by implicit biases, and scientists do not exist in an objectivity vacuum. Subconscious biases in scientists’ choices about which species to study may beget distorted knowledge bases and stagnant paradigms. Disparities in biological knowledge can result from bias in study species selection within a cycle of policymaking, funding, and publication, all subject to implicit biases. Here, we show that ornithological research in the USA and Canada is biased toward birds with greater aesthetic salience and those with larger breeding ranges and ranges that encompass more universities. We quantified components of aesthetic salience (e.g., color, pattern/contrast, body size) of 293 passerines and near-passerines based on empirically documented human visual preferences and investigated whether these components were associated with research effort. We also quantified each species’ breeding range size and the number of universities within that range. Accounting for phylogenetic relatedness, we found that these metrics of aesthetics, familiarity, and accessibility combined to explain 45% of the variation in the number of published papers about each species from 1965–2020. On average, birds in the top 10% of aesthetic salience were studied 3.0X more than birds in the bottom 10%, and publication numbers were predicted most strongly by color and pattern components of aesthetic salience. Birds in the top 10% of breeding range size and university abundance were studied 3.8X and 3.5X more often than species in the bottom 10% of those categories, respectively. Species listed as Endangered and those featured on journal covers have greater aesthetic salience scores than other species. We discuss how these biases may influence perceived relative value of species with respect to culture and conservation. The disparities in empirical knowledge we describe here perpetuate a positive feedback loop, thus widening the gap between the avian “haves” and “have-nots”, with some questions answered repeatedly while potentially critical discoveries are left undiscovered.

> “All animals are equal, but some animals are more equal than others.” —George Orwell, *Animal Farm* (1945)

## Introduction

Human aesthetic values and preferences have been shaped by evolution, culture, and experience. Our color preferences in particular are thought to be adaptive because humans are typically drawn to colors associated with favorable objects or conditions that confer fitness benefits (Palmer and Schloss 2010). For example, humans are drawn to blues, likely because blues are associated with clean water and clear sky (Palmer and Schloss 2010), and we tend to associate greens with healthy nature and calmness and yellows with happiness (Clarke and Costall 2008). On average, humans also tend to be drawn to familiar (e.g., the mere-exposure effect [Zajonc 1968], the familiarity-attraction hypothesis [Reis et al. 2011]), and emotionally positive things (Ishizu et al. 2023). When humans view an object, multiple systems in the brain interact, resulting in an “aesthetic experience” (Chatterjee and Vartanian 2014). Reward centers in the brain are activated when an object is perceived as aesthetically pleasing (Erk et al. 2002), resulting in inherent, subconscious judgments and biases toward “beautiful” objects that elicit pleasure (i.e., positive reinforcement; Prum 2017, Ishizu et al. 2023). Importantly, humans have been described as the “world’s greatest evolutionary force” (Palumbi 2001), and, thus, given our natural biases, what are the consequences for nonhuman species?

Human value judgments about nonhuman species (e.g., plants and nonhuman animals) have contributed to global “biological annihilation” (Ceballos et al. 2017) and thus necessitated the entire concept of conservation as a “crisis” discipline (Soulé 1985). For example, overexploitation and perceived species value in the global wildlife trade (Vall-llosera and Cassey 2017, Senior et al. 2022) have manufactured a species salience hierarchy. One of the primary postulates of conservation biology is that all species have intrinsic value (Soulé 1985); however, such value may not be equally distributed among species (Macdonald et al. 2015). Humans make value-based judgments about which species are “worthy” of conserving and thus where limited conservation dollars are spent (Brambilla et al. 2013, Curtin and Papworth 2020, Adamo et al. 2022), decisions which are often influenced by aesthetic salience (i.e., human interest in or attention to species based on their visual appeal) and charismatic phenotypes (Lorimer 2007, Veríssimo et al. 2014, Williams et al. 2023). In general, humans favor species perceived as aesthetically pleasing, such as larger animals (Berti et al. 2020, Curtin and Papworth 2020), those with bright (Langlois et al. 2022) or multiple colors (Curtin and Papworth 2020), and those that share similarities with humans (Gunnthorsdottir 2001, Batt 2009). Beyond visual appeal, humans also favor species that are geographically familiar (Correia et al. 2016, Mittermeier et al. 2020). Recent “culturomics” studies have analyzed shifts in word frequencies, such as query patterns in search engines (e.g., searches for species names in Google; Ladle et al. 2016, Schuetz and Johnston 2019), to directly quantify species salience, further illuminating human organismal preferences on a larger scale. These subjective preferences can have tangible consequences for conservation, policy, and research (Wilson et al. 2007, Martín-López et al. 2009), and thus for our understanding of biology as a whole.

Birds are the most colorful of the land vertebrates to the human eye despite our inability to see the full range of color or complexity of hues that birds display and perceive (Eaton 2005, Stoddard and Prum 2011). Relative to many other taxa, birds are well studied (Bonnet et al. 2002, Clark and May 2002)—especially in North America and Europe (Ducatez and Lefebvre 2014)—and receive considerable conservation support, but some birds may be disproportionately favored in conservation and research due to human biases. On average, humans tend to prefer bird species that are larger (Correia et al. 2016, Schuetz and Johnston 2019), more colorful (Lišková and Frynta 2013, Echeverri et al. 2020), and boldly patterned (Frynta et al. 2010, Vall-llosera and Cassey 2017, Garnett et al. 2018, Schuetz and Johnston 2019, Andrade et al. 2022). Specifically, evidence suggests that people prefer birds with blue, yellow, and green plumage (Lišková and Frynta 2013, Senior et al. 2022), lighter plumage (Lišková and Frynta 2013), and birds with crests (Echeverri et al. 2020; but see Schuetz and Johnston 2019). Birdwatchers disproportionately seek out and report colorful, boldly patterned species compared to their drab counterparts (Echeverri et al. 2020, Stoudt et al. 2022). Similarly, a study of online photographs of bird plumage aberrations by Zbyryt et al. (2021) indicated a collection bias towards larger bird species with larger populations adjacent to human settlements. Taken together, these studies broadly point to human preferences towards colorful, boldly patterned birds (i.e., aesthetically salient) that are nearby and familiar. Functional magnetic resonance imaging (fMRI) shows that birdwatchers and ornithologists perceive and recognize birds similarly to human faces (Gauthier et al. 2000), especially in locally familiar species (Wing et al. 2022), which may enhance the potential for bias in study species selection and research effort. Importantly, aesthetic salience refers to visual prominence or noticeability, and its antonym is unremarkable, not ugly: with respect to garnering attention from humans, unremarkable is likely worse than ugly (e.g., smooth-headed blobfish [*Psychrolutes marcidus*], naked mole rat [*Heterocephalus glaber*], California condor [*Gymnogyps californianus*]).

Cultural values can influence conservation, policy, and scientific research funding (Wilson et al. 2007), hence shaping research topics and priorities, thereby enabling a complex positive feedback loop (Martín-López et al. 2009). For example, species that are societally salient (often due to aesthetic appeal) may be conservation and/or research priorities, thus generating additional research effort that may inform policy, conservation status decisions and/or wasted conservation dollars (Buxton et al. 2021), and future scientific interest (Martín-López et al. 2009). Scientists do not exist in a vacuum of pure objectivity and are influenced by cultural values (Wilson et al. 2007, Jarić et al. 2019). There are many examples of public attitudes and biases in conservation preferences, but fewer that assess potential bias in scientists and their research choices (e.g., study species). Some studies have quantified taxonomic biases in research effort/attention to identify factors that contribute to study species selection (Jarić et al. 2015, Fleming and Bateman 2016, da Silva et al. 2020, Guedes et al. 2023). Organisms that are more broadly distributed (Jarić et al. 2015, Fleming and Bateman 2016, Yarwood et al. 2019) and/or more geographically proximal to humans (Guedes et al. 2023) often receive greater research effort and attention than rarer species found farther away from human settlements. Few papers quantify research effort disparities in terms of aesthetic preferences toward species traits (e.g., Bellwood et al. 2020, Adamo et al. 2021). Aesthetically salient fish (Bellwood et al. 2020) and plants (Adamo et al. 2021) garner greater research effort and attention among scientists. However, inequity in research effort relative to overall aesthetic salience of birds has not been addressed.

Here, we identify and quantify a disparity in research effort associated with human visual perception, geographic accessibility, and familiarity within 293 North American birds. We do not attempt to define “beauty”, assign relative value to species, or create a priority ranking of birds. Instead, we systematically quantify parameters empirically demonstrated to vary in human visual preferences of birds (e.g., color, lightness, body size, pattern/contrast) to test whether variation in aesthetic salience among species is associated with variation in the number of peer-reviewed publications featuring those species. In addition, we use breeding range sizes and the number of institutions of higher education within each breeding range as proxies for species’ familiarity and accessibility, respectively, to researchers in the continental United States and Canada. We quantify research effort from 1965–2020 using an intensive bibliometric approach and hypothesize that birds with greater aesthetic salience, larger breeding ranges, and ranges that encompass more universities have historically been more well studied than their small, drably plumaged, and distant-from-humans counterparts. We also investigate differences in the aesthetic salience of species listed as Endangered and those featured on covers of scientific journals. We discuss how these biases may preclude important discoveries, influence perceived relative value of species, contribute to decisions about naming species, and influence choices of species exploited for fundraising.

## Materials and Methods

### Species selection

We studied 293 passerine and near-passerine species in the continental United States and Canada and accounted for changes in taxonomy and nomenclature. For additional details, see *Supplementary material, Species selection and nomenclature*.

### Bibliometrics

Using an intensive bibliometric approach, we quantified 1965–2020 research effort for each species. We used the number of publications (i.e., a count of the number of peer-reviewed book chapters, journal articles, etc. [hereafter, papers]) indexed by Web of Science (www.webofscience.com) as a proxy for research effort. To search Web of Science, we created Boolean search strings that accounted for changes in species taxonomy and nomenclature (see above) and downloaded each species’ raw output as a csv file. We subsequently conducted a systematic, qualitative assessment of each species’ search output to further constrain our dataset to include only papers in which the species was a primary focal species (i.e., one of up to three focal species; *Supplementary material, Bibliometric data*). The final paper count for each species was recorded and used as the response variable in models.

### Scoring species’ aesthetic salience

We created a system to assign each species an Aesthetic Salience Score (ASS), a cumulative value that incorporates documented human visual preferences towards birds (Lišková and Frynta 2013, Lišková et al. 2015, Curtin and Papworth 2020, Echeverri et al. 2020). We quantified 7 characteristics (*Supplementary material, Table S1*) including color score, body mass (a proxy for size), lightness, contrast/pattern, iridescence, presence and size of a crest, and presence of other extraordinary features (e.g., tail streamers). We performed our assessments using color illustrations of each species obtained from the Sibley Birds v2 mobile phone application (https://www.sibleyguides.com/product/sibley-birds-v2-app/) to standardize imagery and avoid variability in quality, lighting, clarity, and editing associated with photographs. Detailed methodology and justification is in *Supplementary material, Aesthetic salience score (ASS)*.

### Estimating species familiarity and accessibility: range size and institutions of higher learning

To test for familiarity and accessibility bias in research effort on North American birds, we determined each species’ breeding range size and the number of institutions of higher learning (hereafter, “universities”) within each breeding range. We calculated species’ breeding range sizes (km^2^) in ArcMap (v10.7.1; Environmental Systems Research Initiative [ESRI]) using shapefiles obtained from BirdLife International (2021). To determine the number of universities within each species breeding range, we compiled a map of 4-year colleges and universities in the US and Canada. We overlaid breeding range maps with the map of institutions and extracted the number of colleges and universities within each species’ range in ArcMap. Detailed methodology for estimating species’ familiarity and accessibility is available in *Supplementary material, Range size and institutions of higher learning*.

### Phylogeny and statistical analysis

We obtained a phylogenetic tree from birdtree.org (Jetz et al. 2012). We pruned the ultrametric Hackett tree (birdtree.org/subsets) to include the 293 species represented in our study and downloaded 1000 trees to account for phylogenetic uncertainty. We then used the program TreeAnnotator (v2.6.2) to generate a single maximum clade credibility tree. Final node heights were estimated using the common ancestor heights.

Prior to conducting any analyses, we evaluated the correlations among the continuous predictor variables to determine whether there was evidence of multicollinearity. We used the function “vif” in the *car* package in R (R Core Team 2023) to identify predictor variables with high correlations by calculating variance inflation factors. All variance inflation factor values were below 5, suggesting no evidence of multicollinearity. Thus, no highly correlated predictor variables were included in the model.

We used a phylogenetic comparative approach to quantify the variables that predicted the number of publications for each bird species in our data. Because the response variable (number of publications) consisted of counts, we performed a poisson regression. All continuous traits were standardized using the “standardize” function in the *effectsize* package in R (Ben-Shachar et al. 2020). Standardizing the predictor variables avoids the problems of traits that differ in scale. In addition, using standardized variables enhances the interpretation of the regression analysis.

We applied a Markov Chain, Monte Carlo generalized linear mixed model (MCMCglmm) to assess the contribution of each predictor variable to the number of publications using the *MCMCglmm* package in R (Hadfield 2010). We ran the MCMCglmm including phylogeny as a random effect. The model was run for 400,000 iterations with a burnin of 10,000 and thinning interval of 100. As suggested by Hadfield (2010), we used uninformative priors. We ran each model with four chains and checked for convergence with the Gelman-Rubin diagnostic (Brooks and Gelman 1998). We also calculated the phylogenetic heritability, which may be used as an estimate of phylogenetic signal. We first ran a model that included the number of publications as the response variable and overall ASS, breeding range size of each species (km^2^), number of universities within the breeding range of the species, and body mass. We next assessed the contribution of the aesthetic components to the overall ASS using a MCMCglmm phylogenetic model. Based on the results from this analysis we performed an additional MCMCglmm analysis with only the significant predictors of ASS. We estimated the marginal means of the predictor variables using the *ggeffects* package in R (Lüdecke 2018).

## Results

We considered 293 species of passerines and near-passerines whose breeding ranges were entirely or substantially within the continental US and/or Canada (see *Supplementary material, Species selection and nomenclature*). In total, we downloaded and considered publication information from 27,254 papers from 1965– 2020 in our literature search, only 50% (13,620/27,254) of which we retained in our dataset based on our filtering criteria (see *Supplementary material, Bibliometric data*). The median number of papers published per bird species was 19 (IQR = 40, *xˉ* = 46 ± 79 SD, range 0–597; Fig.1). The 5 species with the most papers were tree swallow (*Tachycineta bicolor*; *n* = 597), red-winged blackbird (*Agelaius phoeniceus*; *n* = 499), song sparrow (*Melospiza melodia*; *n* = 469), white-crowned sparrow (*Zonotrichia leucophrys*; *n* = 465), and black-capped chickadee (*Poecile atricapillus*; *n* = 444). Ten species (3%) were the primary focus of zero papers (e.g., black-chinned sparrow [*Spizella atrogularis*] and crissal thrasher [*Toxostoma crissale*]).

**Figure 1.**
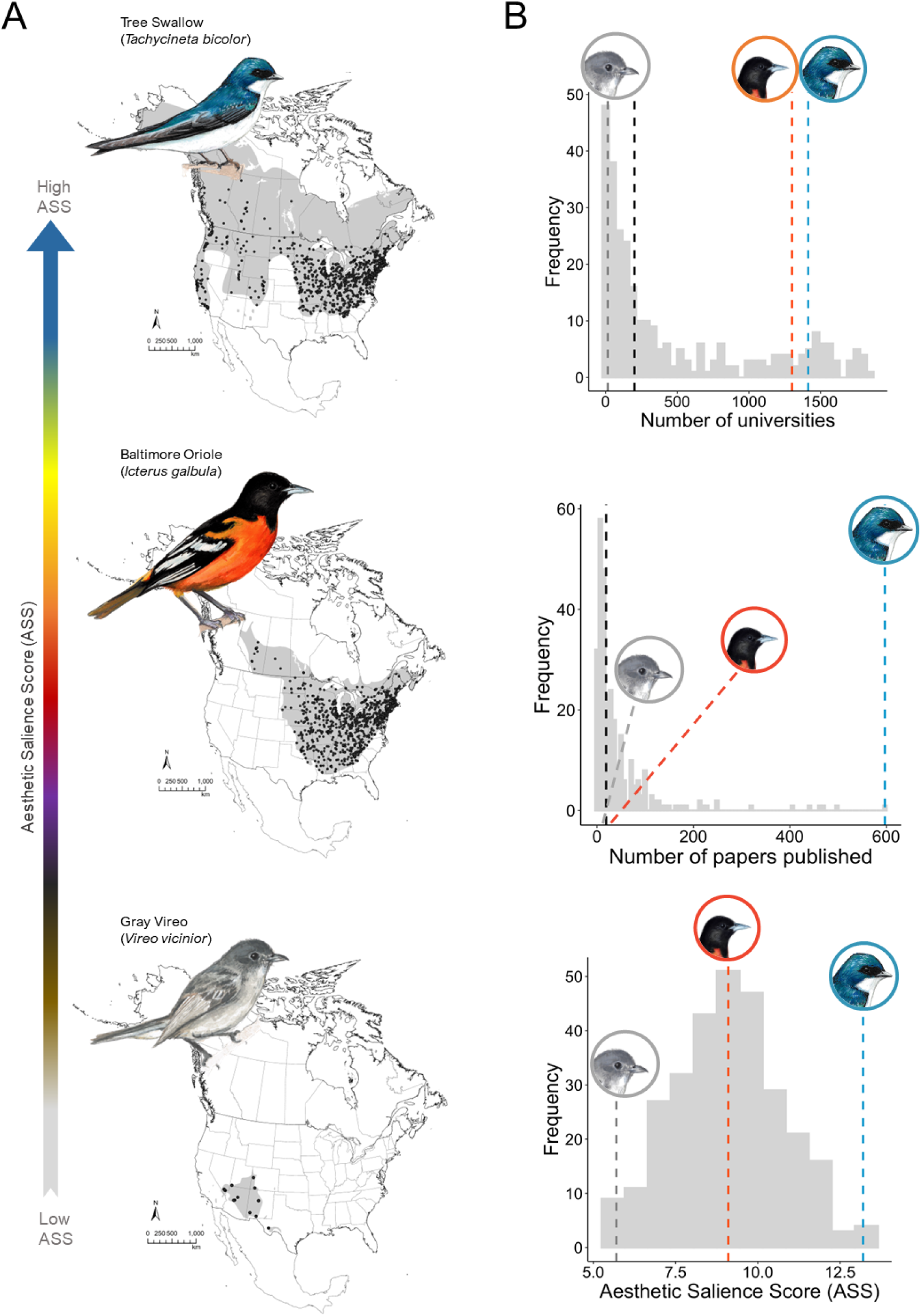
A) Examples of species with high (tree swallow; *Tachycineta bicolor*), medium (Baltimore oriole; *Icterus galbula*), and low (gray vireo; *Vireo vicinior*) Aesthetic Salience Scores (ASS). Species’ breeding range maps (gray fill) were accessed from BirdLife International (2021). Black dots depict individual universities within species ranges. B) Histograms on the right indicate data spread of the number of universities within species’ breeding ranges, number of papers published per species, and Aesthetic Salience Scores (ASS). Black dashed lines in histograms denote medians across the dataset. Gray, orange, and blue dashed lines denote where the three species occur in the data distribution. Note that Baltimore oriole ASS was equal to median ASS; thus, the orange line in the ASS histogram also represents the dataset median. Species illustrations by Silas E. Fischer.

Values for the Aesthetic Salience Score (ASS), a cumulative metric incorporating color, contrast, lightness, body mass, and other visual characteristics (see *Supplementary material, Aesthetic salience score [ASS]* and *Table S1*), ranged from 5.5–13.2 (xˉ = 9.1 ± 1.6 SD; *Supplementary material, Fig. S1*). The species with the greatest ASS of 13.2 were bohemian waxwing (*Bombycilla garrulus*) and tree swallow, whereas black swift (*Cypseloides niger*) and chimney swift (*Chaetura pelagica*) had the lowest ASS of 5.5.

### Predictors of number of publications

We evaluated the contributions of ASS, breeding range size, and number of universities in the breeding range (hereafter, “number of universities”) to explain the variation in number of publications. The model revealed significant effects for ASS, breeding range size, and the number of universities (*Supplementary material, Table S2*). Thus, species with a high value for ASS, a large breeding range, and a large number of universities had a greater number of publications (Fig. 2). Specifically, species in the top 10% of ASS were studied 3.0 times more than birds in the bottom 10%, on average (Fig. 2). Species in the top 10% of breeding range size and university abundance were studied on average 3.8 times and 3.5 times more often than species in the bottom 10% of those categories, respectively (Fig. 2). A MCMCglmm mixed model analysis assessing the contribution of the seven aesthetic characteristics to the number of publications revealed that only two (i.e., color score and contrast/pattern; posterior means = 0.29 and 0.23, respectively; p < 0.001; *Supplementary material, Table S3*) were significant predictors. We found no significant contribution of lightness, body mass, iridescence, crest, or extraordinary features in explaining variation in the number of publications. In fact, we analyzed the relative influence of the main color of each species, color score, contrast/pattern, lightness, iridescence, presence of a crest, and other extraordinary features to the overall ASS. A surprising result was the non-significant effect of the main color on ASS. The remaining variables were all significant contributors to overall ASS (*Supplementary material, Table S3*). We then analyzed the influence of color score, contrast/pattern, iridescence, presence of a crest, and other extraordinary features on the number of publications. We also included breeding range size, number of universities, and body mass as covariates. The analysis revealed that inclusion of the aesthetic features iridescence, crest, lightness, and extraordinary features did not predict the number of publications (Table 1). The breeding range size and the number of universities had the largest effect sizes followed by color score and contrast/pattern (Table 1).

**Figure 2.**
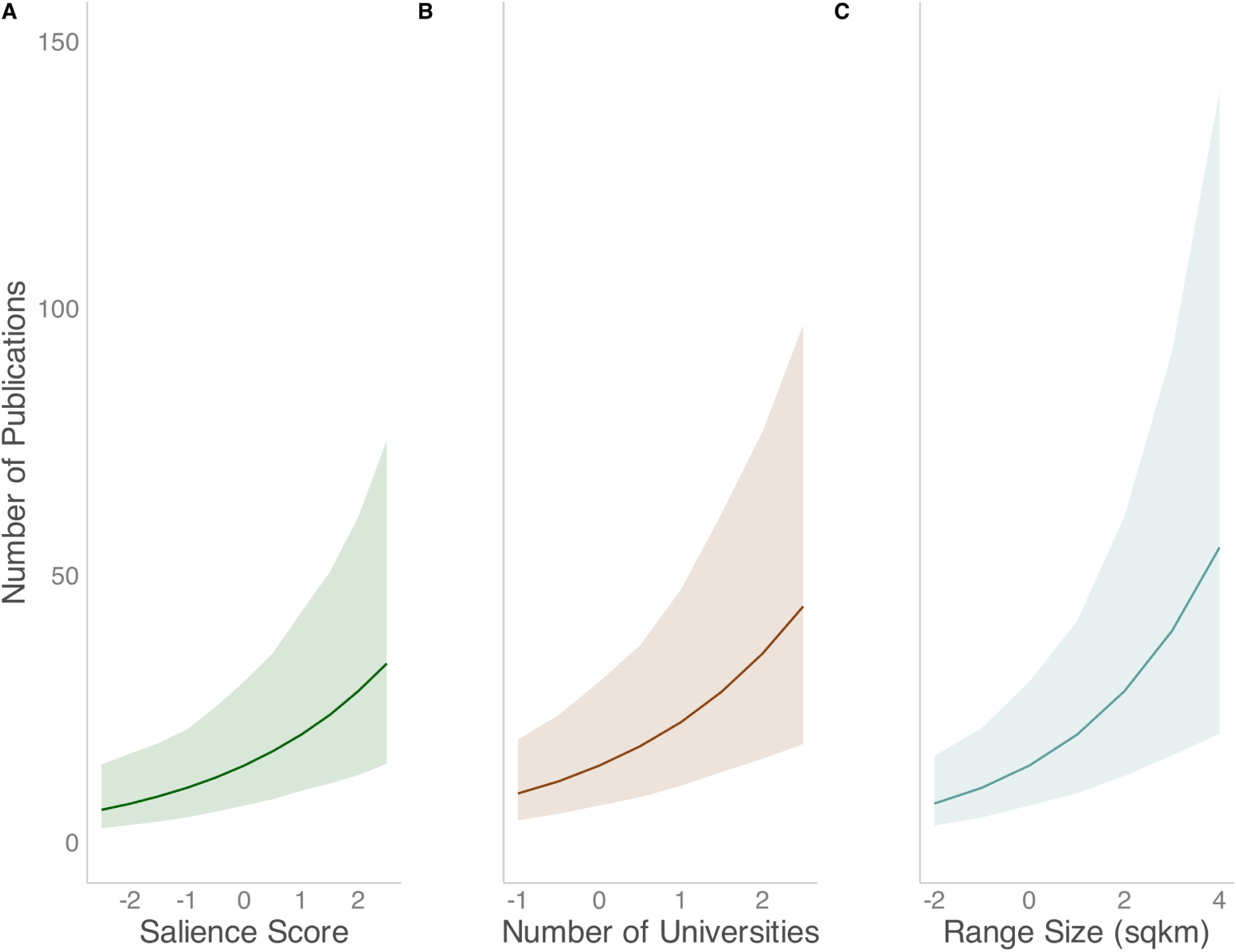
Marginal mean number of publications for the three significant predictor variables: A) Aesthetic salience score, B) Number of universities within species’ breeding ranges, and C) Breeding range size (km^2^). Note that the predictor variables are scaled to a mean of 0 and SD = 1.

**Table 1.**
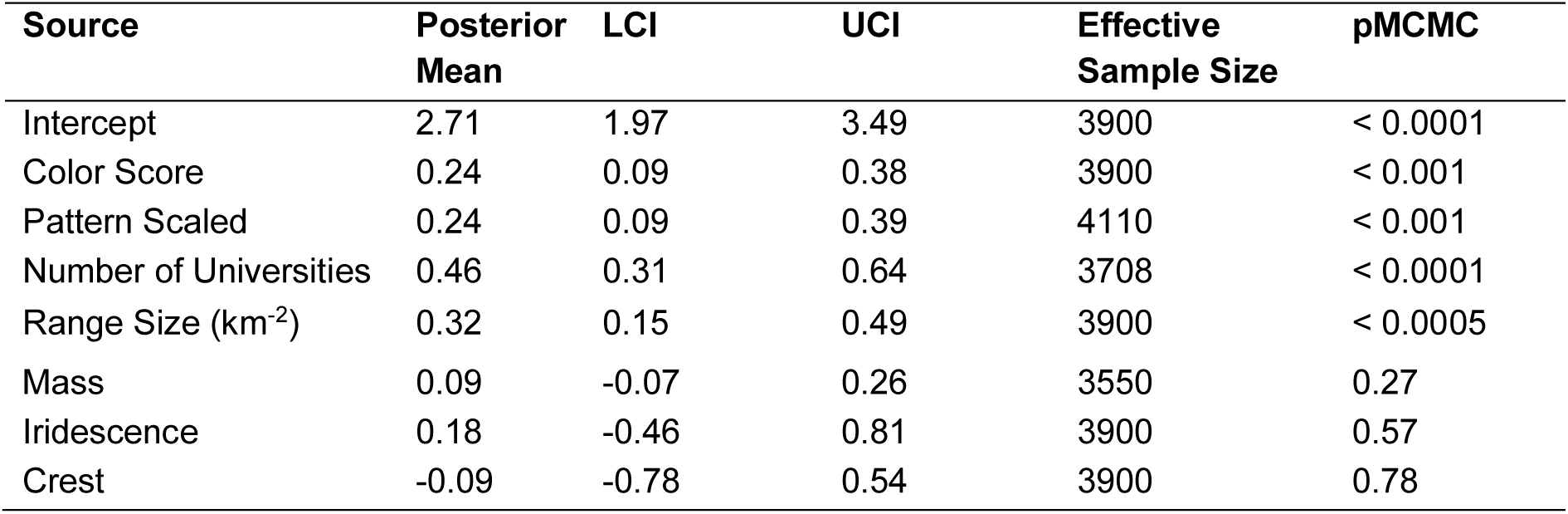
MCMCglmm phylogenetic models determining the contribution of significant predictors of Aesthetic Salience Score (ASS; see *Supplementary material, Table S3*), breeding range size, and number of universities within breeding ranges to the number of publications about each 293 species from 1965–2020. The continuous variables were scaled to a mean of 0 and standard deviation of 1.0. The phylogenetic heritability (equivalent to Pagel’s λ) = 0.92. Values presented are the posterior means, 95% lower (LCI) and upper (UCI) credible intervals, effective sample sizes, and p-values (pMCMC).

### Endangered species

Of the 293 species of interest, 9 were listed as Endangered in the US and 13 were listed as Endangered in Canada (2 of those were listed in both countries; *Supplementary material, Endangered species*). Aesthetic salience scores (ASS) for species listed as Endangered in the US (xˉ_ASS_ = 9.98 ± 0.30 SE) were greater than ASS for species not listed as Endangered in the US or Canada (xˉ_ASS_ = 9.04 ± 0.10 SE; *t* = 0.99, *df* = 275, *P* = 0.01). The ASS for species listed as Endangered in Canada (xˉ_ASS_ = 9.45 ± 0.42 SE) were relatively similar to ASS for species not listed as Endangered in the US or Canada (xˉ_ASS_ = 9.04 ± 0.10 SE; *t* = 0.72, *df* = 279, *P* = 0.36). The difference between countries appeared to be driven by Canada recognizing the lowest ASS bird in our dataset (black swift, ASS = 5.5) as Endangered. Excluding the black swift, the other 12 species listed as Endangered in Canada have greater ASS than species not listed as Endangered in either country (*t* = 0.98, *df* = 278, *P* = 0.03).

### Journal covers

We reviewed the covers (1965–2021) of 29 peer-reviewed scientific journals (*Supplementary material, Journal covers*) and found 143 instances of 94 of our species of interest featured on a cover. Birds that were featured on covers had greater ASS (xˉ_COVER-ASS_ = 9.48 ± 0.15 SE) compared to those not featured on covers (xˉ_NON-COVER-ASS_ = 8.90 ± 0.12 SE; *t* = 2.9, *df* = 291, *P* = 0.005).

## Discussion

We found that there has been greater research effort on aesthetically pleasing birds that occur in larger breeding ranges that encompass more universities, indicating a significant bias in the focus of ornithological research in the USA and Canada. Science plays a pivotal role in a cycle that dictates public interest/awareness, policy, conservation status, conservation and management decisions/actions, and research funding disparities (Martín-López et al. 2009), thus feeding a snowball of bias accumulation by iteratively producing greater research effort, attention, and citations to a select few species because of their appearance and not necessarily their potential to progress science. In a highly competitive research environment where success is often measured by money coming in and publications going out, scientists may have little choice but to perpetuate this problem, even if they are aware of it, rather than risk career advancement. Such an aesthetic bias feedback loop emulates the “Matthew effect” (i.e., the rich get richer and the poor get poorer; Zhang et al. 2015) and further divides the organismal “haves” and “have-nots” in a world with limited research and conservation resources and accelerating biodiversity loss. Furthermore, many forms of research bias, including bias toward charismatic species, can lead to wasted conservation efforts (Buxton et al. 2021).

Combined, our variables representing aesthetic salience, familiarity, and accessibility accounted for 45% of the variation in published research among bird species in the US and Canada. It was not our objective to explain all of the variation in research effort in all birds, which could also be associated with charismatic behaviors, vocalizations, cognition, habitat associations, value as game species, model species, and many other factors. For example, across all birds, migratory species are studied 10-times more often than resident species (Ducatez and Lefebvre 2014), a factor that we did not consider here. However, we note that most of these other variables are not free from entanglement with biases in human visual perception (e.g., migratory species tend to be more colorful and have lighter-colored feathers than non-migrants; Delhey et al. 2021). We found that body mass was not associated with research effort. However, we note that our study was constrained to passerines and near-passerines (i.e. a narrow range of body masses) and therefore is not necessarily in contradiction with studies detecting mass-related biases when comparing a wider range of taxa and body masses (e.g., shrews [Soricidae] to elephants [Elephantidae] and hummingbirds [Trochilidae] to ostriches [*Struthio* sp.]; Ducatez and Lefebvre 2014, Tam et al. 2022; Guedes et al. 2023). We also found no association between research effort and the presence or size of a bird’s crest, consistent with birdwatchers’ preferences in Schuetz and Johnston (2019) but not in Echeverri et al. (2022). We note that both of those studies included a broader range of bird species and investigated interests of birdwatchers, most of whom are not ornithologists and may exhibit greater aesthetic biases as a group. We also found no association between research effort and plumage lightness, iridescent plumage, or extraordinary features like tail streamers, all of which might be expected to contribute to aesthetic appeal to humans.

Declines of North American passerines have been associated with many factors including climate change, habitat loss, exotic predators, and collisions with man-made objects (Loss et al. 2015), but not with human visual preference to our knowledge. Therefore, the exceptionally great ASS of the birds with Endangered status may reflect aesthetic bias in the process and support for petitioning and listing species. Even the extinct ivory-billed woodpecker (*Campephilus principalis*), last definitively observed in 1944 and therefore not studied in the wild during our period of interest, continues to hold Endangered status in the US, with >20M $US spent over the past two decades on predictably failed searches (Troy and Jones 2023). We included only extant species in our study, but we speculate that the prestige bias behind much of the ivory-billed woodpecker ignominy (Troy and Jones 2023) is likely bolstered by aesthetic bias, as the species would score in the 97th percentile (ASS = 12.2) of our metric. It is difficult to imagine a similar financial boom, media bonanza, and cult-like public following would develop around researchers at a regional college legitimately rediscovering a thought-to-be extinct small, drab passerine.

Our finding that both breeding range size and number of universities within a species’ range were independently significant predictors of research effort in US and Canada birds corroborates previous studies assessing geographic bias in research on birds (Ducatez and Lefebvre 2014, Murray et al. 2015, Yarwood et al. 2019) and other taxa (e.g., non-avian reptiles [Guedes et al. 2023], mammals [Fleming and Bateman 2016], and plants [Adamo et al. 2021]). This research effort disparity suggests that many ornithologists may be selecting study species on logistical convenience, which likely results from limited funding. Beyond research effort, plant species with larger range sizes receive more conservation funding, potentially because a greater number of researchers within larger ranges can access funding (Adamo et al. 2022). Even some bird species with relatively great ASS, such as Scott’s Oriole (*Icterus parisorum*; ASS = 9.7) have received very little research attention, likely due to a relatively small breeding range size and relatively great distance from universities. Thus, small, drab birds with small ranges farther from universities, such as many birds in the Southwestern US (e.g., gray vireo [*Vireo vicinior*; ASS=5.7], gray flycatcher [*Empidonax wrightii*; ASS=7.6], bushtit [*Psaltriparus minimus*; ASS=5.6]), are rarely represented in the scientific literature. Furthermore, Ducatez and Lefebvre (2014) demonstrated that island bird species are less studied than those that occur on the “mainland.” Many birds in the Southwestern USA occur on “sky islands” (McCormack et al. 2009), which may contribute to reduced research effort despite such species technically occurring on the “mainland.” What exciting and important discoveries are we omitting from our knowledge base by excluding drab birds in even more remote locations around the world while ever more studies focus on, for example, American redstart (*Setophaga ruticilla*; ASS = 9.9) habitat associations?

Although time-consuming, the experience of filtering through the Web of Science output compels us to caution against using raw exported numbers of “hits” from literature search engines as comparative measures of the quantity of research focused on given entities, as is common in studies of research effort (e.g., Ibáñez-Álamo et al. 2017, da Silva et al. 2020, Guedes et al. 2023). The presence of a species name in a paper, or even its abstract, does not ensure a study was focused on that species, and some species are more likely than others to be named in studies in which they are not a primary focus. As two examples, we found that only 10% (17/162) of papers produced by our search for alder flycatcher (*Empidonax alnorum*) were actually papers focused on that species, compared to 82% (72/89) for prothonotary warbler (*Protonotaria citrea*), a difference that is attributable primarily to the former being noted more often than the latter in general community studies. We note that Web of Science indexed many but not all peer-reviewed outlets, and thus some papers were not counted here (e.g., we found some of our own studies were not included). However, we have no reason to believe these exclusions were taxonomically biased, and thus they should not impact our results.

That birds on journal covers had greater ASS than those not featured on covers is unsurprising regardless of any bias in research rates because journal editors and publishers likely select cover images they believe will catch the attention of readers. In other words, human bias toward fancy birds is not only recognized, but also strategically exploited for marketing purposes. Papers that are featured in journal cover images often garner greater Altmetric scores (www.altmetric.com; Kong and Wang 2020) and more citations relative to non-cover papers in the same journal (Wang et al. 2017), and thus can perpetuate the taxonomic bias feedback loop. This bias recognition is similar to selecting aesthetically pleasing species as “poster children” or “flagships” for conservation efforts and fund-raising campaigns (Garnett et al. 2018). But these terms are not biological concepts (Veríssimo et al. 2011) and care should be taken that they are not exploited to justify management or conservation actions empirically demonstrated to negatively impact the purported beneficiary species (e.g., Gross 2005, Streby et al. 2018).

Human inability to perceive color differences that birds see between males and females of many species (Eaton 2005), and our aesthetic biases when we can differentiate sexual dichromatism, might both contribute to female birds being relatively rarely studied (Gaines et al. 2020) and underrepresented in museum collections (Cooper et al. 2019). Additionally, human color preferences can be reflected in the names we attribute to birds (Stoddard and Prum 2011) and may subsequently hinder some species in terms of research potential. Would the gray vireo be more well studied if it had been given a more exciting name like vivacious vireo, reflecting its lively nature rather than its relative colorlessness? Would the American dipper (*Cinclus mexicanus*) be less well studied if it had been named the dusky ditch-dipper? In our dataset, birds with common names starting with “black”, “gray”, “brown”, or “plain” (the most common colors of birds; Delhey et al. 2023) were studied approximately half as often as all other birds. As committees debate the renaming of birds currently assigned honorific or eponymous common names (Driver and Bond 2021), we encourage them to acknowledge human biases and consider the potential advantages and disadvantages of naming species based on human visual perception. As an extreme example (not a recommendation), the renaming of Bewick’s wren (*Thryomanes bewickii*) to rambunctious wren versus brown wren could influence the potential for future research on the species.

Aesthetic biases may also contribute to discrepancies in conservation and management efforts between closely related species. For example, the golden-winged warbler (*Vermivora chrysoptera*) and blue-winged warbler (*V. cyanoptera*) are typically described as a hybridizing species complex (e.g., Buehler et al. 2007). Both “species” have experienced numerically stable global population trends in the 21st century (Ziolkowski et al. 2022). Yet the northern, more aesthetically salient golden-winged warbler (ASS = 9.6) has been studied substantially more (*n* = 101 papers) than the more southerly, less aesthetically salient blue-winged warbler (ASS = 8.6; *n* = 30 papers), with many of the blue-winged warbler papers being focused on their association with, and presumed negative impact on, golden-winged warblers.

Interbreeding with the blue-winged warbler is often assumed to be a cause of regional declines in Golden-winged Warblers (e.g., Confer et al. 2020), leading some to suggest conservation efforts should aim to keep blue-winged warblers away from golden-winged warblers (e.g., Confer et al. 2020) and that geographic segregation would be a desirable outcome of climate change (Hightower et al. 2023). This pseudo-conflict is inconsistent with evidence that genetic introgression is symmetrical (Shapiro et al. 2004) and has largely genetically homogenized the system (Toews et al. 2016) but is consistent with the implicit bias toward segregation in some human demographics (Anicich et al. 2021). Despite nearly indistinguishable genomes (Toews et al. 2016), we are not aware of any calls to lump golden-winged and blue-winged warblers into one species, and we speculate that this is at least partly because their minor differences are in genes associated with plumage color (Toews et al 2016). Neglect, and even vilification, of less aesthetically salient birds is thought-provoking considering human tendency to equate “beauty” with moral “goodness” and thus make overall judgments based on attractiveness (i.e., the “halo effect”; Batres and Shiramizu 2022). Darwin (1859) said, “I have been struck with the fact, that if any animal or plant in a state of nature be highly useful to man, or from any cause closely attract his attention, varieties of it will almost universally be found recorded. These varieties, moreover, will often be ranked by some authors as species (p 61–62) … I was much struck how entirely vague and arbitrary is the distinction between species and varieties” (p 60). We postulate that golden-winged and blue-winged warblers would be considered one species from their original description if their minor genetic differences were associated with other traits (e.g., gut morphology, nestling growth rates, saliva production) not subject to human aesthetic bias. Indeed, species in our dataset that have been lumped at any time during 1965–2020 had a below average mean ASS (8.1) while those that have been split have 16% greater mean ASS (9.4). Therefore, it is possible that many multi-species complexes are currently considered a single species partly or solely because they are all drab and/or mostly indistinguishable with human visual perception. If true, then not only are aesthetically salient birds studied more often in general, but the problem may be compounded by aesthetic bias in attributing species status in the first place.

When Charles Darwin embarked on his famous expedition aboard the H.M.S. *Beagle*, he was enthralled by, for example, the “beautifully coloured” Planarians (Darwin 1839, p. 27) of Brazil and the “brilliantly coloured” birds of Maldonado (p. 40) but unimpressed by the drab organisms inhabiting the Galápagos Islands (e.g., “…The insects…are small-sized and dull-coloured…All the plants have a wretched, weedy appearance, and I did not see one beautiful flower.”; p 381). Darwin remarked that “…none of the birds are brilliantly coloured, as might have been expected in an equatorial district” (1839, p 381). Hence, Darwin initially may not have been overly interested in the small, drab-plumaged Galápagos “finches” (Thraupidae; Sulloway 1982, 1984; Lack 1953), which were first described by a previous explorer as “…not remarkable for their novelty or beauty” (Lack 1947, p 8–9). Indeed, Darwin did not even correctly record the localities of the representative “finch” specimens he collected (Sulloway 1982). Thus, his interest in the “finches” was largely retrospective (Lack 1947, Sulloway 1982). Had Darwin not studied this system, now a textbook example of natural selection and adaptive radiation, due to his initial biases against the “…drab sameness of their plumage” (Weiner 1994, p. 434), where would the biological sciences be today? What would we have missed out on without Darwin’s insights—without studying species that might not be as visually appealing to the human eye? With the growing understanding that awareness is the first step in overcoming implicit biases in many areas of human life (e.g., Lee 2017), it is our sincere hope that a concerted effort will emerge in ornithology to discover what we’ve been missing as we have neglected drab and distant birds.

## Acknowledgments

We are grateful for initial input on study conceptualization and design from B.E. Carpenter, E. Zeigler, M. Rizk, and T.L. Spanbauer. We thank J. Refsnider, J. Zubcevic, and C.E. Nemes for helpful comments on the manuscript prior to submission. D.B. Miles was supported by NSF grant DEB 1950636.

## Author Contributions

S.E.F., J.G.O., A.M.L., and H.M.S. designed research; S.E.F., J.G.O., A.M.L., and H.M.S. performed research; D.B.M. analyzed data; S.E.F. and D.B.M. created figures, and S.E.F., J.G.O., A.M.L., D.B.M., and H.M.S. wrote the paper.

## Supplementary Material

### Species selection and nomenclature

We focused our study on passerines (Passeriformes) and “near-passerines” in North America. We followed standardized common and scientific names published in The American Ornithological Society’s (AOS; formerly American Ornithologists’ Union [AOU]) species checklist of birds in North and Central America, using all volumes and supplements from the 5th Edition (1957) through the 61st Supplement (Chesser et al. 2020). We used the following criteria to determine which species to include for analysis: a) only North American species from the orders Passeriformes, Cuculiformes, Apodiformes, Coraciiformes, and Piciformes (i.e., passerines and “near-passerines”), b) only species that have considerable breeding populations within the mainland United States (US) and/or Canada (i.e., excluding species for which the range is almost entirely outside of these countries), and c) only native species. We consulted various sources that describe species breeding distributions (e.g., field guides) to determine which species to include. We acknowledge that “considerable breeding population” is somewhat subjective but a) we needed some type of delineation, b) the goal was to include only species for which more than vagrants or a small number of breeding pairs occupied the area of interest, and c) there is no reason to believe this delineation introduced any bias to our dataset. Introduced species such as house sparrow (*Passer domesticus*) and European starling (*Sturnus vulgaris*), game species (e.g., Columbiformes), and brood parasites (i.e., brown-headed cowbird [*Molothrus ater*]) were excluded. We excluded the parasitic brown-headed cowbird because its unique breeding strategy elicits intensive research effort compared to species that provide parental care (Ducatez and Lefebvre 2014).

For each species, we determined whether it was “split” into multiple species (i.e., multiple species that were previously recognized as one species; *n* = 8), or whether multiple species were “lumped” into one (i.e., one species that was previously recognized as multiple species; *n* = 3) within the first half of our period of interest (i.e., up to 1993; see below for midpoint criterion). Additionally, 120 species required synonyms to be included in our bibliometric search (see *Supplementary material, Bibliometric data*) because of a) a common and/or scientific name change, b) a name change from lumping or splitting after 1993, or c) international spelling differences (e.g., “grey” and “gray”). Considering name synonyms minimizes the potential for omission of relevant publications in our bibliometric search (Correia et al. 2018). To determine if and when nomenclature and/or taxonomic changes occurred, we used the AOS/AOU checklist supplements published throughout the period of interest. If a species’ scientific or common name was changed during the period of interest, but the species was not lumped or split, we included all of that species’ common and scientific names as synonyms in our literature search and we identified the species in our dataset by its name as of 31 December 2020. For species that were lumped or split between 1965–2020, we included them as per their classification during the majority of the period. That is, if a bird was split into multiple species, we included them as one species (i.e., the pre-split name) if the split occurred more than halfway through the period of interest (i.e., 1993 or later); if a split occurred prior to 1993, we used the post-split species names and we used information from each pre-split paper to determine which of those species was studied. For example, yellow-bellied sapsucker (*Sphyrapicus varius*) split in 1983 to yellow-bellied sapsucker and red-breasted sapsucker (*S. ruber*), and then in 1998 also split off red-naped sapsucker (*S. nuchalis*). Using the 1993 midpoint criterion, we retained yellow-bellied sapsucker and red-breasted sapsucker, whereas red-naped sapsucker was considered a synonym for yellow-bellied sapsucker but not a separate species for our analysis. We treated “lumped” species similarly, whereby we included multiple species if the lumping occurred 1993 or later, and included only the single, post-lump species if it was lumped prior to 1993. In one case, (i.e., gilded flicker [*Colaptes chrysoides*] and northern flicker [*C. auratus*]), two species were lumped prior to 1993 and re-split after 1993, and were therefore separate species for the majority of our period of interest; thus we considered these as separate species. Our goal was to make consistent, repeatable rules for determining how to include species and we’re not sorry if your favorite species was not included the way you would prefer. In total, we included 293 species in our analyses.

### Bibliometric data

We obtained bibliometric data from the Web of Science (www.webofscience.com) during February 2022 to quantify research effort (i.e., a count of the number of peer-reviewed book chapters, journal articles, etc. [hereafter, papers]) for each of the 293 species. We searched all papers published between 1 January 1965 and 31 December 2020 in Web of Science using the standardized common and scientific names, as described above. A csv file with pertinent publication data (e.g., title [TI], topic [TS], abstract [AB], author [AU], etc.) or each species was constructed by using a Boolean search string that included:

1) Topic (TS) “Common Name” or
2) Abstract (AB) “Common Name” or
3) Topic (TS) “Genus species” or
4) Abstract (AB) “Genus species” or
5) Any synonyms also searched in Topic and Abstract with OR’s and “ “

For example, the search for papers on the plain titmouse (*Parus inornatus*; a species that was split into oak titmouse [*Baeolophus inornatus*] and juniper titmouse [*Baeolophus ridgwayi*] in 1996) included the following string: TS= “Plain Titmouse” OR AB= “Plain Titmouse” OR TS= “Parus inornatus” OR AB= “*Parus inornatus*” OR TS= “Oak Titmouse” OR AB= “Oak Titmouse” OR TS= “Juniper Titmouse” OR AB= “Juniper Titmouse” OR TS= “*Baeolophus inornatus*” OR AB= “*Baeolophus inornatus*” OR TS= “*Baeolophus ridgwayi*” OR AB= “*Baeolophus ridgwayi*”.

Notably, search options in Web of Science changed after we collected these bibliometric data, and a search of “topic” now includes the title, key words, and abstract, which would reduce the string options required to replicate our search.

We limited our searches to only species’ names that were located in the topic or abstract to constrain our initial bibliometric data to those papers in which the target species was included in the study. We subsequently conducted a qualitative assessment of each species’ Web of Science search output to further constrain our dataset to include only papers in which the species was a primary focal species of a study. This assessment involved reading the title and abstract of each paper and, when necessary, reviewing the published paper to exclude additional papers in which the species was not the primary focus. We excluded results that were not full research publications (e.g., published conference abstracts); we excluded papers that had nothing to do with birds and for which we could not determine why Web of Science included them; we excluded papers that mentioned a species in the abstract for comparative purposes but for which the species was not a focus of the study; and we excluded studies focused on a different species elsewhere in the world with similar or identical common names (e.g., the ovenbirds [Furnariidae] of South and Central America and multiple species of yellow warblers in Africa). We limited inclusion of papers in our dataset to all papers that were clearly single-species studies and we also included: a) hybridization studies, b) multi-species studies with 3 or fewer bird species, c) ecto– and endo-parasite studies with 3 or fewer bird species, d) vagrant records outside of a species’ expected distribution, and e) non-ecology studies (e.g., kinematics of hummingbirds compared to vehicles; and papers in journals such as “*Bioinspiration & Biomimetics*” or “*Nutrition Journal*”).

A primary purpose of this filtering was to isolate the number of papers that focused on each species and to avoid biasing our publication counts toward species that commonly appear in broader community studies. For example, many community point-count survey studies in eastern North America list ovenbird (*Seiurus aurocapilla*) in the abstract simply because it was among the most common species recorded in a community, but not because ovenbirds were a focus of the paper. Through this process, we further refined our dataset to studies that took place in the mainland US and Canada and studies outside that area (e.g. a non-breeding ground study) where we could confirm an author affiliation in the US or Canada because we had to somehow limit the scope or we would add too many additional known and unknown variables. We acknowledge that our search for papers was not exhaustive as we have used only a single bibliographic database and focused our search on topics and abstracts; however, we believe that, following our extensive vetting of the initial output, any remaining bias was similar across species and thus unlikely to affect patterns in relative publication numbers.

### Aesthetic salience score (ASS)

We created a system to assign each species an Aesthetic Salience Score (ASS) by quantifying the likely visual appeal to humans of the 293 bird species of interest. Our ASS system is based upon human perception of color rather than birds’ perception because human perception drives the visual appeal of birds to people. We first obtained a color plate illustration for each species from the Sibley Birds v2 mobile phone application (https://www.sibleyguides.com/product/sibley-birds-v2-app/) to standardize visual comparisons among species images and limit potential biases in image color, lighting, contrast, etc. (i.e., issues with comparing photographs). For nearly all species (*n* = 288), we used profile-view illustrations of adult males in breeding plumage. When profile-view illustrations were not available (*n* = 4; all of which were swift [Apodidae] species), we used in-flight illustrations. When profile-view illustration was available for the female but not the male (*n* = 1; northern flicker), we used the female illustration because the sexes appear very similar in that species. When multiple profile-view illustrations were available (e.g., illustrating regional plumage variants), we randomly selected from the available illustrations (*n* = 89).

We quantified total ASS for each species’ image using a systematic, repeatable process we created for assigning cumulative values among seven categories associated with human visual preference in birds (Lišková and Frynta 2013, Lišková et al. 2015, Curtin and Papworth 2020; Table S1). We weighted the scores for each category such that they would contribute differently to total ASS based on relative importance inferred from previous studies of aesthetic appeal of birds and other imagery to humans. These features, in order of estimated importance to human aesthetic perception, included color score, contrast/pattern, body mass (a proxy for size), lightness, iridescence, presence and size of a crest, and presence of other extraordinary features (e.g., tail streamers; Table S1). For color, we referenced the Sibley images to determine each species’ plumage colors (i.e., black, blue, brown, gray, green, orange, purple, red, white, and yellow) as a group (i.e., the authors came to consensus on which colors were present). We intentionally did not use computer programs or other automated methods to distinguish between colors because we were specifically interested in colors readily differentiated by human perception, as humans perceive color categorically (Bird et al. 2014). We did not attempt to differentiate among minor differences in color hues. Each bird was assigned a cumulative score for all colors present, such that blue, yellow, and green were given greater value (1.5 pt each) than intermediate preference colors including red, orange, and purple (1 pt each), and the less preferred black, white, gray, and brown (0.5 pt each). For example, if a bird was black (0.5 pt), blue (1.5 pt), and white (0.5 pt), it received a color score of 2.5 pts.

We quantified pattern/contrast using the Contrast CoVAnalysis macro in the PAT-GEOM v1.0.1 plugin (Chan et al. 2018) in ImageJ (Schneider et al. 2012), which produces a measure of the coefficient of variation of pixel tonal values within a selected portion (i.e., the bird) of an image. We chose to use PAT-GEOM in ImageJ to quantify contrast because, compared to human ability to discretize colors (Bird et al. 2014), contrast is more difficult to manually quantify (Chan et al. 2018). We multiplied each contrast score by 2000 to scale the mean to 3.2 and max to 7.5 and scale the variable to contribute similar weight as the color score variable to total ASS. We obtained mean body mass (g) values for each species from Dunning (2007), from which we used species mean values when sexes were not differentiated, and we used mean male mass when reported separately. The distribution of masses was non-normal with many small birds and relatively few heavier birds; therefore, we natural-log transformed the mass variable. We then divided the transformed mass values by two to scale the variable to contribute approximately half as much weight as the color variable to total ASS.

Lightness values were calculated in Adobe Photoshop Elements 2021 (Adobe Inc.) by selecting the body of each bird in the Sibley image, excluding background and feet, and viewing the tonal histogram which displays and summarizes the tonal range of all pixels within the selected portion of an image. Tonal values are unitless and range from 0 (darkest) to 255 (lightest) representing brightness regardless of color. We used the median tonal value for each bird because many histograms did not follow normal distributions. We divided tonal values by 70 to scale the mean to 1.7 and max to 3.3 and scale the lightness variable to contribute approximately half as much weight as the color score variable to total ASS. Lastly, we added 1 pt to each species’ ASS for the presence of each additional visually appealing characteristic, including iridescent plumage (i.e., humans prefer shiny things likely because of innate need for water; Meert et al. 2014), presence and size of a crest (Echeverri et al. 2020; but see Schuetz and Johnston 2019), and other extraordinary features (e.g., tail streamers, notably bright or contrasting eye color, waxy feather tips) that may be associated with visual appeal to humans (Frynta et al. 2010). Iridescence, presence and size of a crest, and other extraordinary features were assigned and agreed upon by the authors.

### Range size and institutions of higher learning

We calculated 292 species’ breeding range sizes (km^2^) in ArcMap (v10.7.1; Environmental Systems Research Initiative [ESRI]) using shapefiles obtained from BirdLife International (2021). In one case, BirdLife International did not have range map shapefiles (i.e., hoary redpoll [*Acanthis hornemanni*]); thus, we referenced a current range map (https://www.audubon.org/field-guide/bird/hoary-redpoll) and manually drew a polygon of the breeding range in ArcMap. To determine US institutions of higher learning, we started with the USGS college and university dataset, which is composed of all United States Post-Secondary Education facilities as defined by the Integrated Post-Secondary Education System (https://www.sciencebase.gov/catalog/item/4f4e4acee4b07f02db67fb39). A comparable GIS layer for Canadian colleges and universities proved difficult to obtain despite consultation with very friendly and supportive Canadian government employees. Therefore, we used Google Maps as a reference, searched “college” and “university”, scrolled across the map of Canada, and manually added points to our GIS layer for all entities that could be confirmed to be 4-yr colleges and universities. These points were accurate to within ∼1 km, and not to exact address locations, which was adequate for the scale of our analysis. Across both countries, we included only doctoral/research universities, masters colleges and universities, and baccalaureate colleges (i.e., we excluded community colleges, technical/vocational schools, etc., because they tend to have less active research programs). We did not screen every institution to determine whether ornithology research has been conducted there within the period of interest. Rather, within the US institution GIS layer, we manually screened the attributes table and removed all institutions that were unlikely to host ornithology or ecology research of any kind. For example, we excluded seminaries, independent medical schools, law schools, art, music, and culinary institutes, business schools, and all 73 University of Phoenix locations. In addition, we excluded duplicate locations (e.g., multiple campuses across a city or off-campus buildings identified independently in the shapefile) within a city for an institution, such that each college or university was represented by one point.

### Endangered species

To compare aesthetic salience scores of species listed as Endangered at the federal level with those not listed as Endangered, we retrieved US and Canada federal conservation status information for each species following the United States Fish and Wildlife Service’s (USFWS) Environmental Conservation Online System (ECOS; USFWS 2019) list under the Endangered Species Act (ESA) and the Environment and Climate Change Canada (ECCC) list under the Species At Risk Act (SARA; Canada). We included a species as Endangered if the entire species, one or more subspecies, or a distinct population segment was listed as Endangered in either or both countries during our period of interest (i.e., even if the species was recently delisted, e.g., Kirtland’s warbler [*Setophaga kirtlandii*] in the US). We compared ASS of species listed as Endangered in each country (*n* _US_ = 9; *n*_Canada_ = 13) to ASS of species not listed as Endangered in either country (*n* = 268). The total of these three counts does not match the total birds studied because 2 species were listed as Endangered in both countries.

### Journal covers

We reviewed cover images from 29 relevant peer-reviewed journals that publish cover images (i.e., mainly photographs but some illustrations)—or did so during some portion or our period of interest—associated with papers in each issue to compare the ASS of “cover birds” to birds that were not featured on covers. We counted the number of instances each of our species of interest was represented in a cover image of a peer-reviewed journal. Journals we reviewed with cover images included *Ardea*, *Auk* (renamed *Auk: Ornithological Advances*, then renamed *Ornithology*), *Behavioral Ecology*, *Bird Conservation International*, *Condor* (renamed *Condor*: *Ornithological Applications*, then renamed *Ornithological Applications*), *Conservation Biology*, *Current Biology*, *Diversity and Distributions*, *Ecography*, *Ecological Applications*, *Ecology*, *Ecology Letters*, *Functional Ecology*, *Frontiers in Ecology and the Environment*, *Global Change Biology*, *Journal of Animal Ecology*, *Journal of Applied Ecology*, *Journal of Avian Biology*, *Journal of Wildlife Management*, *Landscape Ecology*, *Methods in Ecology and Evolution*, *Nature*, *OIKOS*, *Ornithological Monographs*, *Proceedings of the National Academy of Sciences USA*, *Proceedings of the Royal Society of London B*, *Science*, *Studies in Avian Biology*, and *Wildlife Society Bulletin*. We excluded covers that displayed collages of more than 3 bird species.

---

**Supplementary material, Table S1.** Aesthetic salience characteristics quantified and investigated as potential predictors of number of publications (1965-2020).

| Characteristic | Values | Reference(s) | Significant predictor of no. publications? |
| --- | --- | --- | --- |
| Color score |  |  |  |
| (those distinctly present on a bird's plumage, scored by consensus among the authors) | 1.5 each: Blue, Green, Yellow | Lišková and Frynta (2013), Echeverri et al. (2020), Senior et al. (2022) | Yes |
|  | 1.0 each: Red, Orange, Purple |  |  |
|  | 0.5 each: Black, Brown, Gray, White |  |  |
| Contrast/Pattern |  |  |  |
| (coefficient of variation of pixel tonal values measured with Contrast CoVAnalysis macro in the PAT-GEOM v1.0.1 plugin [Chan et al. 2018] in ImageJ [Schneider et al. 2012]; value divided by 70 to scale contrast value to contribute approximately the same as Color score to ASS) | Mean 3.2; Range 1.0–7.5 | Frynta et al. (2010), Vall-Ilosera and Cassey (2017), Garnett et al. (2018), Schuetz and Johnston (2019), Andrade et al. (2022) | Yes |
| Body mass |  |  |  |
| (mean mass in grams from Dunning [2007], ln transformed, then divided by 2 to scale body mass value such that it contributed half as much as Color score to ASS) | Mean 1.6; Range 0.5–3.4 | Correia et al. (2016), Schuetz and Johnston (2019) | No |
| Lightness |  |  |  |
| (median tonal value from Histogram in Adobe Photoshop Elements 2021; multiplied by 2000 to scale Lightness value such that it contributed half as much as Color score to ASS) | Mean 1.7; Range 0.4–3.3 | Lišková and Frynta (2013) | No |
| Iridescence |  |  |  |
| (absence, presence) | 0, 1 | Meert et al. (2014) | No |
| Crest |  |  |  |
| (absence, presence small, presence sizeable; scored by consensus among the authors) | 0, 0.5, 1 | Echeverri et al. (2020) | No |
| Other extraordinary features |  |  |  |
| (presence of at least one; e.g., tail streamers, waxy tips of feathers, striking eye color) | 0, 1 | Frynta et al. (2010) | No |
| Total ASS | Mean 9.1; Range 5.5–13.2 |  |  |

**Supplementary material, Table S2.** MCMCglmm phylogenetic models determining the contribution of Aesthetic Salience Score (ASS) to the number of publications about each 293 species from 1965–2020. We included the number of universities within each breeding range, the breeding range size, and body mass of each species as covariates. The continuous variables were scaled to a mean of 0 and standard deviation of 1.0. The phylogenetic heritability (equivalent to Pagel’s λ) = 0.92. Values presented are the posterior means, 95% lower (LCI) and upper (UCI) credible intervals, effective sample sizes, and p-values (pMCMC).

| Source | Posterior<br>Mean | LCI | UCI | Effective<br>Sample Size | pMCMC |
| --- | --- | --- | --- | --- | --- |
| Intercept | 2.64 | 1.89 | 3.37 | 4700 | < 0.001 |
| **ASS** | **0.34** | **0.19** | **0.49** | **3900** | **< 0.001** |
| **Number of**<br>**Universities** | **0.45** | **0.28** | **0.62** | **3900** | **< 0.001** |
| **Range Size (km<sup>2</sup>)** | **0.33** | **0.16** | **0.49** | **3900** | **< 0.001** |
| Mass (g) | 0.05 | -0.11 | 0.21 | 3900 | 0.51 |

**Supplementary material, Table S3.** MCMCglmm analysis identifying the component traits associated with Aesthetic Salience Score (ASS). The phylogenetic heritability (equivalent to Pagel’s λ) = 0.99. Values presented are the posterior means, 95% lower (LCI) and upper (UCI) credible intervals, effective sample sizes, and p-values (pMCMC).

| Source | Posterior Mean | LCI | UCI | Effective Sample Size | pMCMC |
| --- | --- | --- | --- | --- | --- |
| Intercept | -0.03 | -0.28 | 0.18 | 3900 | 0.77 |
| *Main Color* |  |  |  |  |  |
| Blue | 0.03 | -0.07 | 0.12 | 3900 | 0.59 |
| Brown | 0.04 | -0.02 | 0.09 | 3900 | 0.22 |
| Gray | 0.04 | -0.02 | 0.09 | 3900 | 0.19 |
| Green | -0.03 | -0.11 | 0.05 | 3900 | 0.52 |
| Orange | 0.03 | -0.07 | 0.12 | 4193 | 0.53 |
| Pink | -0.13 | -0.33 | 0.09 | 3900 | 0.23 |
| Red | -0.02 | -0.12 | 0.10 | 3900 | 0.78 |
| White | 0.00 | -0.06 | 0.06 | 3904 | 0.97 |
| Yellow | 0.02 | -0.05 | 0.09 | 3900 | 0.64 |
| Color Score | 0.59 | 0.58 | 0.61 | 3900 | < 0.0001 |
| Pattern Scaled | 0.68 | 0.66 | 0.71 | 3900 | < 0.0001 |
| Lightness Scaled | 0.28 | 0.25 | 0.30 | 3900 | < 0.0001 |
| Iridescence | 0.61 | 0.50 | 0.71 | 3900 | < 0.0001 |
| Crest | 0.75 | 0.63 | 0.87 | 3871 | < 0.0001 |
| Extra Feature | 0.75 | 0.68 | 0.83 | 3900 | < 0.0001 |

**Supplementary material, Figure S1.**
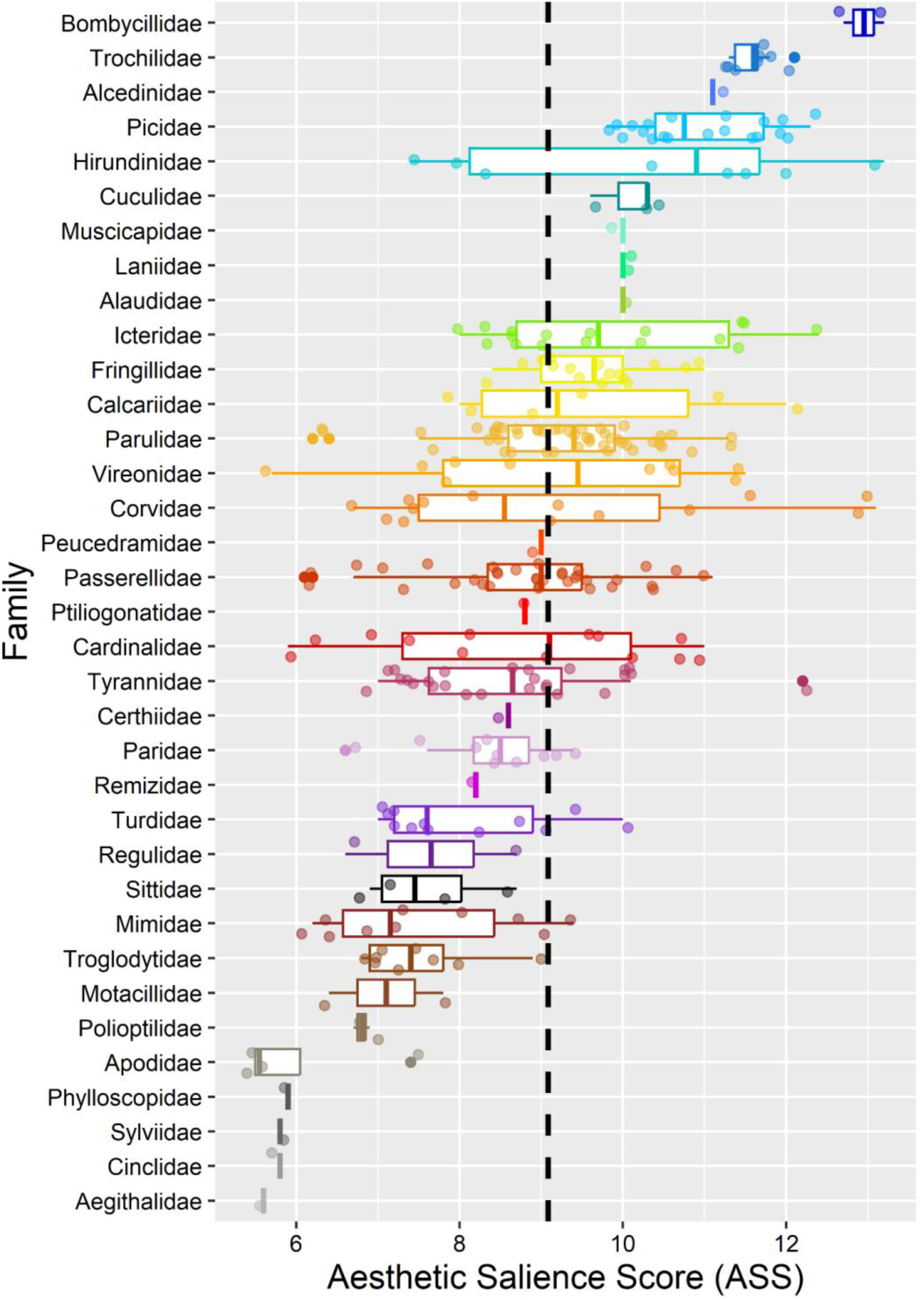
Aesthetic Salience Scores (ASS) of taxonomic families in order of highest (top; Bombycillidae) to lowest (bottom; Aegithalidae) score. Thick interior bars depict median, boxes span first and third quartiles (inter-quartile range [IQR]), whiskers depict minimum and maximum, and jittered dots denote extreme values. The vertical dashed black line indicates mean ASS across all families (*xˉ*_ASS_ = 9.1).

## Literature Cited

M. Adamo et al., Plant scientists’ research attention is skewed towards colorful, conspicuous, and broadly distributed flowers. Nat Plants 7, 574–578 (2021).

M. Adamo et al., Dimension and impact of biases in funding for species and habitat conservation. Biol Conserv 272, 109636 (2022).

R. Andrade et al., Species traits explain public perceptions of human–bird interactions. Ecol Appl 32, e2676 (2022).

E.M. Anicich, J.M. Jachimowicz, M.R. Osborne, L.T. Phillips, Structuring local environments to avoid racial diversity: Anxiety drives Whites’ geographical and institutional self-segregation preferences, J Exp Soc Psychol 95, 104117 (2021).

M.D. Baiz et al., Genomic and plumage variation in *Vermivora* hybrids. Auk 137, 1– 14 (2020).

C. Batres, V. Shiramizu, Examining the “attractiveness halo effect” across cultures, Curr Psychol 2022, 1–5 (2022).

S. Batt, Human attitudes towards animals in relation to species similarity to humans: a multivariate approach. Biosci Horiz 2, 180–190 (2009).

D.R. Bellwood, C.H. Hemington, S.B. Tebbett, Subconscious biases in coral reef fish studies. Biosci 70, 621–627 (2020).

M.S. Ben-Shachar, D. Lüdecke, D. Makowski, effectsize: Estimation of effect size indices and standardized parameters. J Open Source Softw 5, 2815 (2020).

E. Berti, S. Monsarrat, M. Munk, S. Jarvie, J-C. Svenning, Body size is a good proxy for vertebrate charisma. Biol Conserv 251, 108790 (2020).

BirdLife International and Handbook of the Birds of the World. 2021. Bird species distribution maps of the world. Version 2021.1. Available at http://datazone.birdlife.org/species/requestdis.

X. Bonnet, R. Shine, O. Lourdais, Taxonomic chauvinism. Trends Ecol Evol 17, 1–3 (2002).

M. Brambilla, M. Gustin, C. Celada, Species appeal predicts conservation status. Biol Conserv 160, 209–213 (2013).

S.P. Brooks, A. Gelman, General methods for monitoring convergence of iterative simulations. J Comput Graph Stat 7, 434–455 (1998).

D.A. Buehler et al., Status and conservation priorities of Golden-winged Warbler (*Vermivora chrysoptera*) in North America. Auk 124, 1439–1445 (2007).

R.T. Buxton, et al., Avoiding wasted research resources in conservation science. Conserv Sci Prac 3, e329 (2021).

G. Ceballos, P.R. Ehrlich, R. Dirzo, Biological annihilation via the ongoing sixth mass extinction signaled by vertebrate population losses and declines. Proc Natl Acad Sci USA 114, E6089–E6096 (2017).

A. Chatterjee, O. Vartanian, Neuroaesthetics. Trends Cogn Sci 18, 370–375 (2014).

J.A. Clark, R.M. May, Taxonomic bias in conservation science. Science 297, 191– 192 (2002).

T. Clarke, A. Costall, The emotional connotations of color: a qualitative investigation. Color Res Appl 33, 406–410 (2008).

J.L. Confer et al., Implications for evolutionary trends from the pairing frequencies among Golden-winged and Blue-winged Warblers and their hybrids. Ecol Evol 10, 10633–10644 (2020).

N. Cooper et al., Sex biases in bird and mammal natural history collections. Proc Royal Soc B 286, 20192025 (2019).

R.A. Correia, P.R. Jepson, A.C.M. Malhado, R.J. Ladle, Familiarity breeds content: assessing bird species popularity with culturomics. PeerJ 4, e1728 (2016).

P. Curtin, S. Papworth, Coloring and size influence preferences for imaginary animals, and can predict actual donations to species-specific conservation charities. Conserv Lett 13, e12723 (2020).

A.F. da Silva et al., Taxonomic bias in amphibian research: Are researchers responding to conservation need? J Nat Conserv 56, 125829 (2020).

C. Darwin, The Voyage of the Beagle (Murray, 1839).

C. Darwin, On the Origin of Species by Means of Natural Selection (Murray, 6th ed., 1859).

K. Delhey, J, Dale, M. Valcu, B. Kempenaers, Migratory birds are lighter coloured. Curr Biol 31, 1511–1512 (2021).

K. Delhey, M. Valcu, C. Muck, J. Dale, B. Kempenaers, Evolutionary predictors of the specific colors of birds. Proc Natl Acad Sci USA 120, e2217692120 (2023).

R.J. Driver, L.J. Bond, Towards redressing inaccurate, offensive and inappropriate common bird names. Ibis 163, 1492–1499 (2021).

S. Ducatez, L. Lefebvre, Patterns of research effort in birds. PLoS One 9, e89955 (2014).

M.D. Eaton, Human vision fails to distinguish widespread sexual dichromatism among sexually “monochromatic” birds. Proc Natl Acad Sci USA 102, 10942– 10946 (2005).

A. Echeverri et al., Can avian functional traits predict cultural ecosystem services? People Nat 2, 138–151 (2020).

S. Erk, M. Spitzer, A.P. Wunderlich, L. Galley, H. Walter, Cultural objects modulate reward circuitry. NeuroReport 13, 2499–2503 (2002).

P.A. Fleming, P.W. Bateman, The good, the bad, and the ugly: which Australian terrestrial mammal species attract most research? Mamm Rev 46, 241–254 (2016).

D. Frynta, S. Lišková, S. Bültmann, H. Burda, Being attractive brings advantages: the case of parrot species in captivity. PLoS One 5, e12568 (2010).

C.D. Gaines, E.M. Rose, K.J. Odom, K.E. Omland, The role of diversity in science: a case study of women advancing female birdsong research. Anim Behav 168, 19–24 (2020).

S.T. Garnett, G.B. Ainsworth, K.K. Zander, Are we choosing the right flagships? The bird species and traits Australians find most attractive. PLoS One 13, e0199253 (2018).

I. Gauthier, P. Skudlarski, J.C. Gore, A.W. Anderson, Expertise for cars and birds recruits brain areas involved in face recognition. Nat Neurosci 3, 191–197 (2000).

L. Gross, Why not the best? How science failed the Florida panther. PLoS Biol 3, Art 333 (2005).

J.J.M. Guedes, M.R. Moura, J.A.F. Diniz-Filho, Species out of sight: elucidating the determinants of research effort in global reptiles. Ecography 2023, e06491 (2023).

A. Gunnthorsdottir, Physical attractiveness of an animal species as a decision factor for its preservation. Anthrozoös 14, 204–215 (2001).

J.D. Hadfield, MCMC methods for multi-response generalized linear mixed models: the MCMCglmm R package. J Stat Softw 33, 1–22 (2010).

J.D. Ibáñez-Álamo, E. Rubio, K. Bitrus Zira, The degree of urbanization of a species affects how intensively it is studied: a global perspective. Front Ecol Evol 5, 41 (2017).

T. Ishizu, T. Srirangarajan, T. Daikoku, “Linking the neural correlates of reward and pleasure to aesthetic evaluations of beauty” in Art and Neurological Disorders, A. Richard, M. Pelowski, B.T. Spee, eds (Humana Press, 2023), pp 215–231.

I. Jarić, J. Knežević-Jarić, J. Gessner, Global effort allocation in marine mammal research indicates geographical, taxonomic and extinction risk-related biases. Mamm Rev 45, 54–62 (2015).

I. Jarić et al., On the overlap between scientific and societal taxonomic attentions — Insights for conservation. Sci Total Environ 648, 772–778 (2019).

W. Jetz, G.H. Thomas, J.B. Joy, K. Hartmann, A.O. Mooers, The global diversity of birds in space and time. Nature 491, 444–448 (2012).

L. Kong, D. Wang, Comparison of citations and attention of cover and non-cover papers. J Informetr 14, 101095 (2020).

D. Lack, Darwin’s Finches (Cambridge University Press, 1947).

D. Lack, Darwin’s finches. Scientific American 188, 66–73 (1953).

R.J. Ladle et al., Conservation culturomics. Front Ecol Environ 14, 269–275 (2016).

J. Langlois et al., The aesthetic value of reef fishes is globally mismatched to their conservation priorities. PLoS Biol 20, e3001640 (2022).

C. Lee, “Awareness as a first step toward overcoming implicit bias” in Enhancing Justice: Reducing Bias 289, S. Redfiled, et al., Eds. GWU Law School Public Law Research Paper No. 2017-56 (2017).

S. Lišková, D. Frynta, What determines bird beauty in human eyes? Anthrozoös 26, 27–41 (2013).

S. Lišková, E. Landová, D. Frynta, Human preferences for colorful birds: vivid colors or pattern? Evol Psychol 13, 339–359 (2015).

J. Lorimer, Nonhuman charisma. Environ Plan D: Soc Space 25, 911–932 (2007).

S.R. Loss, T. Will, P.P. Marra, Direct mortality of birds from anthropogenic causes. Annu Rev Ecol Evol Syst 46, 99–120 (2015).

D. Lüdecke, ggeffects: Tidy data frames of marginal effects from regression models. J Open Source Softw 3, 772 (2018).

E.A. Macdonald, D. Burnham, A.E. Hinks, A.J. Dickman, Y. Malhi, D.W. Macdonald, Conservation inequality and the charismatic cat: *Felis felicis*. Glob Ecol Conserv 3, 851–866 (2015).

B. Martín-López, C. Montes, L. Ramírez, J. Benayas, What drives policy decision-making related to species conservation? Biol Conserv 7, 1370–1380 (2009).

J.E. McCormack, H. Huang, L.L. Knowles, “Sky islands” in Encyclopedia of Islands, R.G. Gillespie, D.A. Clague, eds (University of California Press, 2009), pp. 839–843.

A.J. McKenzie, P.A. Robertson, Which species are we researching and why? A case study of the ecology of British breeding birds. PLoS One 10, e0131004 (2015).

K. Meert, M. Pandelaere, V.M. Patrick, Taking a shine to it: How the preference for glossy stems from an innate need for water. J Consum Psychol 24, 195–206 (2014).

J.C. Mittermeier, U. Roll, T.J. Matthews, R. Correia, R. Grenyer, Birds that are more commonly encountered in the wild attract higher public interest online. Conserv Sci Pract 3, e340 (2021).

H.J. Murray, E.J. Green, D.R. Williams, I.J. Burfield, M. de L. Brooke, Is research effort associated with the conservation status of European bird species? Endanger Species Res 27, 193–206 (2015).

S.E. Palmer, K.B. Schloss, An ecological valence theory of human color preference. Proc Natl Acad Sci USA 107, 8877–8882 (2010).

S.R. Palumbi, Humans as the world’s greatest evolutionary force. Science 293, 1786–1790 (2001).

R.O. Prum, The Evolution of Beauty: How Darwin’s Forgotten Theory of Mate Choice Shapes the Animal World–and Us (Doubleday, 2017).

R Core Team, R: A language and environment for statistical computing. R Foundation for Statistical Computing, Vienna, Austria. https://www.R-project.org/. Version 4.3.1 (2023).

H.T. Reis, M.R. Maniaci, P.A. Caprariello, P.W. Eastwick, E.J. Finkel, Familiarity does indeed promote attraction in live interaction. J Pers and Soc Psychol 101, 557–570 (2011).

J.G. Schuetz, A. Johnston, Characterizing the cultural niches of North American birds. Proc Natl Acad Sci USA 116, 10868–10873 (2019).

R.A. Senior, B.F. Oliveira, J. Dale, B.R. Scheffers, Wildlife trade targets colorful birds and threatens the aesthetic value of nature. Curr Biol 32, 4299–4305 (2022).

L.H. Shapiro, R.A. Canterbury, D.M. Stover, R.C. Fleischer, Reciprocal introgression between Golden-winged Warblers (*Vermivora chrysoptera*) and Blue-winged Warblers (*V. pinus*) in eastern North America. Auk 121, 1019–1030 (2004).

M.J. Silk, S.L. Crowley, A.J. Woodhead, A. Nuno, Considering connections between Hollywood and biodiversity conservation. Conserv Biol 32, 597–606 (2018).

M.E. Soulé, What is conservation biology? Biosci 35, 727–734 (1985).

M.C. Stoddard, R.O. Prum, How colorful are birds? Evolution of the avian plumage color gamut. Behav Ecol 22, 1042–1052 (2011).

S. Stoudt, B.R. Goldstein, P. de Valpine, Identifying engaging bird species and traits with community science observations. Proc Natl Acad Sci USA 119, e2110156119 (2022).

H.M. Streby, G.R. Kramer, S.M. Peterson, S.M., D.E. Andersen, Evaluating outcomes of management targeting the recovery of a migratory songbird of conservation concern. PeerJ 4319 (2018).

F.J. Sulloway, Darwin and his finches: The evolution of a legend. J Hist Biol 15, 1–53 (1982).

F.J. Sulloway, Darwin and the Galapagos. Biol J Linn 21, 29–59 (1984).

J. Tam, M. Lagisz, W. Cornwell, S. Nakagawa, Quantifying research interests in 7,521 mammalian species with h-index: a case study. Gigascience 11, giac074 (2022).

D.P.L. Toews et al., Plumage genes and little else distinguish the genomes of hybridizing warblers. Curr Biol 26, 2313–2318 (2016).

J.R. Troy, C.D. Jones, The ongoing narrative of Ivory-billed Woodpecker rediscovery and support for declaring the species extinct. Ibis 165, 340–351 (2023).

M. Vall-llosera, P. Cassey, Physical attractiveness, constraints to the trade and handling requirements drive the variation in species availability in the Australian cagebird trade. Ecol Econ 131, 407–413 (2017).

D. Veríssimo, D.C. MacMillan, R.J. Smith, Toward a systematic approach for identifying conservation flagships. Conserv Lett 4, 1–8 (2011).

D. Veríssimo et al., Using a systematic approach to select flagship species for bird conservation. Conserv Biol 28, 269–277 (2014).

G. Wang, J. Gregory, X. Cheng, Y. Yao, Cover stories: an emerging aesthetic of prestige science. Public Underst Sci 26, 925–936 (2017).

J. Weiner, The Beak of the Finch (Cambridge University Press, 1994).

T.D. Williams, S. Kreling, L. Stanton, C. Wilkinson, C. Estien, C. Schell, E. Carlen, Of rarity and symbolism: Understanding the human perceptions of charismatic color morphs. Preprint (2023). 10.21203/rs.3.rs-3222187/v1

J.R.U. Wilson, S. Proches, B. Braschler, E.S. Dixon, D.M. Richardson, The (bio)diversity of science reflects the interests of society. Front Ecol Environ 5, 409–414 (2007).

E.A. Wing, F. Burles, J.D. Ryan, A. Gilboa, The structure of prior knowledge enhances memory in experts by reducing interference. Proc Natl Acad Sci USA 119, e2204172119 (2022).

M.R. Yarwood, M.A. Weston, M.R.E. Symonds, Biological determinants of research effort on Australian birds: a comparative analysis. Emu-Austral Ornithology 119, 38–44 (2019).

R.B. Zajonc, Attitudinal effects of mere exposure. J Pers and Soc Psychol 9, 1–27 (1986).

A. Zbyryt, P. Mikula, M. Ciach, F. Morelli, P. Tryjanowski, A large-scale survey of bird plumage colour aberrations reveals a collection bias in Internet-mined photographs. Ibis 163, 566–578 (2021).

H. Zhang, Y. Hu, Y. Zhang, W. Li, Evidence of the Matthew effect in scientific research on mammals in the Chinese First-class National Protected Animals list. Biodivers Conserv 24, 2883–2886 (2015).

D.J. Ziolkowski Jr., M. Lutmerding, V.I. Aponte, M-A.R. Hudson, data from North American Breeding Bird Survey Dataset 1966–2021. Available at 10.5066/P97WAZE5. Deposited 05 July 2022.

## Supplementary Literature Cited

C.M. Bird, S.C. Berens, A.J. Horner, A. Franklin, Categorical encoding of color in the brain. Proc Natl Acad Sci USA 111, 4590–4595 (2014).

I.Z.W. Chan, M. Stevens, P.A. Todd, PAT-GEOM: A software package for the analysis of animal patterns. Methods Ecol Evol 10, 591–600 (2019).

R.T. Chesser et al., Sixty-first supplement to the American Ornithological Society’s check-list of North American birds. Auk 137, 1–24 (2020).

R.A. Correia, et al., Nomenclature instability in species culturomic assessments: Why synonyms matter. Ecol Indic 90, 74–78 (2018).

S. Ducatez, L. Lefebvre, Patterns of research effort in birds. PLoS One 9, e89955 (2014).

J.A. Dunning, CRC Handbook of Avian Body Masses, Second Edition, CRC Press (2007).

D. Frynta, S. Lišková, S. Bültmann, and H. Burda, Being attractive brings advantages: the case of parrot species in captivity. PLoS One 5, e12568 (2010).

S. Lišková, D. Frynta, What determines bird beauty in human eyes? Anthrozoös 26, 27–41 (2013).

C.A. Schneider, W.S. Rasband, K.W. Eliceiri, NIH Image to ImageJ: 25 years of image analysis. Nat Methods 9, 671–675 (2012).

United States Fish and Wildlife Service [USFWS], Environmental conservation online system (ECOS; 2015). https://www.fws.gov/southeast/conservation-tools/environmental-conservation-onlinesystem/.

